# LRRK2 kinase activity is necessary for development and regeneration in *Nematostella vectensis*

**DOI:** 10.1101/2023.11.02.565364

**Authors:** Grace Holmes, Sophie R. Ferguson, Patrick Alfryn Lewis, Karen Echeverri

## Abstract

**Background:** The starlet sea anemone, *Nematostella vectensis*, is an emerging model organism with a high regenerative capacity, which was recently found to possess an orthologue to the human LRRK2 gene (nvLRRK2). The leucine rich repeat kinase 2 (*LRRK2*) gene, when mutated, is the most common cause of inherited Parkinson’s Disease (PD). Its protein product (LRRK2) has implications in a variety of cellular processes, however, the full function of LRRK2 is not well established. Current research is focusing on understanding the function of LRRK2, including both its physiological role as well as its pathobiological underpinnings.

**Methods:** We used bioinformatics to determine the cross-species conservation of LRRK2, then applied drugs targeting the kinase activity of LRRK2 to examine its function in development, homeostasis and regeneration in *Nematostella vectensis*.

**Results:** An *in-silico* characterization and phylogenetic analysis of nvLRRK2 comparing it to human LRRK2 highlighted key conserved motifs and residues. *In vivo* analyses inhibiting the kinase function of this enzyme demonstrated a role of nvLRRK2 in development and regeneration of *N. vectensis*. These findings implicate a developmental role of LRRK2 in *Nematostella*, adding to the expanding knowledge of its physiological function.

**Conclusions:** Our work introduces a new model organism with which to study LRRK biology. We show a necessity for LRRK2 in development and regeneration. Given the short generation time, genetic trackability and in vivo imaging capabilities, this work introduces *Nematostella vectensis* as a new model in which to study genes linked to neurodegenerative diseases such as Parkinson’s.

## Background

The starlet sea anemone *Nematostella vectensis* is a member of the Cnidarian family [1, 2], and is an emerging model for studying development and regeneration due to its ease of maintenance in the lab, sequenced genome and genetic tractability [1, 3–5]. Cnidaria are considered the sister group to the Bilateria, putting them in an excellent position to study the evolutionary trajectory of gene families. Animals from the Cnidarian phylum possess a strikingly similar gene content to humans compared to better studied ecdysozoan models (such as *Drosophila melanogaster*) and possess syntenic genetic loci [6–9]. Of particular interest, *N. vectensis* possesses 4 LRRK genes, one of which being an orthologue to the human *LRRK2* gene [10].

Being composed of two cellular layers, the ectoderm and the endoderm, *Nematostella* have a simple body plan and tissue organization [11–14]. Additionally, *Nematostella* possess a nervous system that takes the form of a nerve net, composed of sensory cells, glandular cells, multipolar ganglion cells and cnidocytes (stinging cells) [4, 15, 16]. Unlike bilaterians, the nervous system of cnidarians is not centralized, and represents an ancestral, simple organization [17, 18]. Nonetheless, several conserved neurogenic pathways and proteins mirroring what is seen in bilaterians highlights their potential usefulness for studying the underlying neurobiology of neurodegenerative diseases like Parkinson’s Disease (PD) [1]. For example, a recent study by Steger et al [8] identified a key upstream regulator of *Nematostella* neuroglandular lineages (composed of neurons, cnidocytes and gland cells): *SoxC,* which has similar roles in bilaterians [19–21]. Prior studies have also demonstrated homologous transcription factors (such as Hox genes) that are involved in neurogenesis, as well as numerous neuropeptides [22, 23]. It is thought that there are 32 potential neuronal cell types in the *Nematostella,* emphasizing the unusual complexity of the nervous system in this organism [24–26].

A key feature of *N. vectensis* is their remarkable ability to regenerate after injury *via* cellular proliferation [27–31]. *Nematostella* can be cut into multiple pieces, and each piece will give rise to a new animal, the exception being that if the foot alone is amputated it cannot regenerate [3, 28, 30]. There has long been an interest in whether or not regeneration reuses the same genes that were developmentally deployed. Recent work from the Rottinger lab has compared gene expression profiles during early development and regeneration and developed a user-friendly website that allows scientists to view the expression of their genes of interest during development *versus* regeneration in *Nematostella vectensis* [3].

LRRK2 is highly studied due to its aforementioned link to PD, with rare autosomal dominant coding mutations as well as more common non-coding variation at the *LRRK2* locus linked to disease risk [32]. The *LRRK2* gene is located on human chromosome 12p11.2-q13.1, and encodes a large 2527 amino acid protein [33–35]. Belonging to the Roco family of Ras-GTPases, LRRK2 is composed of multiple domains (Figure 1) which give rise to multifunctional biology [36, 37].

**Figure 1.**
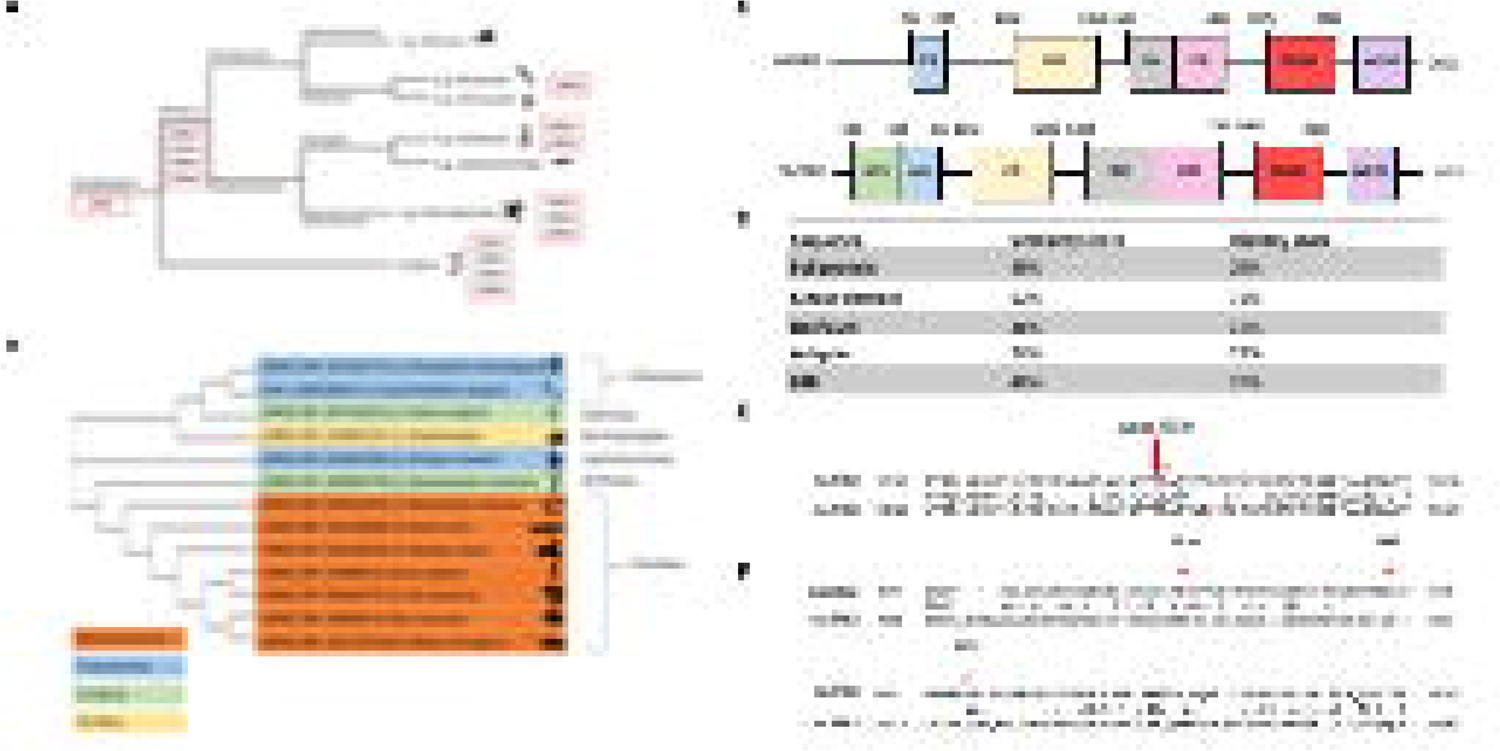
(A) *LRRK* gene orthologues and paralogues across the animalia (B) Phylogenetic sequence analysis of representative LRRK2 proteins (C) Domain structure of nvLRRK2 (NCBI RefSeq: XP_048584778.1) compared to human LRRK2 (hsLRRK2) (NCBI RefSeq: NP_940980.4). (D) Global alignment via Needleman Wunsch algorithm (doi: 10.1016/0022-2836(70)90057-4) illustrating conservation of mutated residues. Arrow indicates most common LRRK2 mutations in PD, G2019S, and I2020T, are conserved in the nvLRRK2 orthologue. (E) Global alignment via Needleman Wunsch algorithm (doi: 10.1016/0022-2836(70)90057-4) illustrating conservation of residues in nvLRRK2 that are commonly phosphorylated in hsLRRK2. Arrow indicates S910 and S935 which are not conserved in nvLRRK2, as well as S973 which is conserved. (F) Percentage similarity and identity score comparing full length nvLRRK2 to hsLRRK2, and individual domains.

Mutations in LRRK2, including N1437H, R1441C, Y1699C, G2019S and I2020T, cluster in the enzymatic domains of the protein and increase LRRK2s kinase activity, with dysregulation of a subset of Rab GTPases a key consequence of this [38]. How this alteration in enzymatic function leads to neurodegeneration is unclear, with studies implicating LRRK2 in a range of cellular processes including lysosomal function, mitochondrial biology and synaptic signaling. The tight association between LRRK2 and Parkinson’s disease has led to a number of in-human clinical trials for both small molecule kinase inhibitors and antisense oligonucleotide gene therapy [39]. Intriguingly, recent work has identified a role for LRRK proteins in stem cell proliferation and tissue remodeling in intestinal enterocytes during whole-body regeneration in the bilaterian freshwater planaria, the *Schmidtea mediterranea* [40] highlighting this as a potential phenotype of interest *Nematostella vectensis*.

In this study we have examined the cross-species conservation of LRRK2 and investigated the functional importance of LRRK kinase activity in *Nematostella* development and regeneration, demonstrating a key role for LRRK2 in these processes.

## Methods

### Animal care

*Nematostella vectensis* were maintained at 17-20°C in Pyrex glass bowls kept in the dark in 15 parts per thousand Instant Ocean [41]. Animals were fed 48-hour old artemia, five times per week. Animals were cleaned a few hours after feeding. Spawning was induced by exposure to light and an increase in temperature to 23-25°C, and embryos were collected immediately after spawning.

### Bioinformatics

#### Evolutionary analysis

To determine the suitability of *N. vectensis* as a model for studying LRRK2-related PD, an evolutionary analysis of LRRK2 genes from different species was carried out. Multiple sequence alignment was carried out using Clustal Omega [42, 43]. The FASTA sequences obtained from National Centre for Biotechnology Information (NCBI) databases [44] were inputted into the program and a phylogenetic tree was generated. The species analyzed were *Nematostella vectensis* (XP_048584778.1), *Homo sapiens* (NP_940980.4), *Mus musculus* (NP_080006.3), *Rattus norvegicus* (NP_001178718.2), *Pan paniscus* (XP_003825774.1), *Danio rerio* (NP_001188385.2), *Xenopus laevis* (XP_018108120.1), *Petromyzon marinus* (XP_032810325.1), *Drosophila melanogaster* (NP_001262772.1), *Caenorhabditis elegans* (BAF48647.1), *Octopus senensis* (XP_029643290.1), i (XP_019853727.1) and the *Hydra vulgaris* (XP_047144213.1). For those species who do not have an exact LRRK2 orthologue, the gene with the greatest percentage similarity was chosen.

#### Domain structure

NCBI reference sequence proteins for human LRRK2 (NP_940980.4) were compared to *Nematostella* LRRK2 (XP_048584778.1). This orthologue was chosen due to having the greatest similarity to the human equivalent. The domain structure was derived from information in the conserved domain databases of NCBI [44]. To compare the similarity between the individual domains of nvLRRK2 and hsLRRK2, the FASTA sequences encoding each domain (based on amino acid positions) were identified and subsequently input into the Needleman-Wunch algorithm via the BLAST database [45–47].

#### Identifying conserved residues

The open reading frames (ORF) of nvLRRK2 and hsLRRK2 were aligned, and the Needleman-Wunsch algorithm was used via the BLAST tool[45, 47] to carry out a pairwise comparison [48]. This provided information including the similarity (amino acids with the same or slightly different side chains) and identity (amino acids in the sequence that are identical). The most common mutated residues in human LRRK2 pertaining to PD were distinguished [49–51], and the corresponding residues in nvLRRK2 were identified. Residues that are conserved have the same amino acid, and those that are not conserved do not. The same technique was used for identifying conservation of phosphorylated residues.

### nvLRRK2 kinase inhibition

To obtain embryos, *N. vectensis* were spawned by exposure to increased temperature and light. The concentration of the kinase inhibitors used for the embryo experiments was 5μm, and for the adult experiments it was 10μm. Some experiments were carried out on Nv-*LWamide* transgenic animals that were expressing mCherry protein their neurons [18]. Embryos were added to a 6-well plate and were incubated in a solution of 5ml sea water and 250μl of each inhibitor. The control embryos were incubated in seawater. For regeneration experiment wild type or transgenic adult *Nematostella* were relaxed in 7.4% MgCl2 (ThermoFisher) for 15 minutes and were subsequently amputated below the pharynx on a plastic petri dish using a sterile no.10 disposable scalpel (World Precision Instruments). The animals were monitored every 2 days and the solutions were changed. Whole animals were fed artemia to ensure any differences were not due to lack of food.

Upon completion of this experiment, the organisms were relaxed in 7.4% MgCl2 (ThermoFisher), and then in 10% MgCl2, 15% MgCl2 and 20% MgCl2. The animals were then fixed in 4% paraformaldehyde (PFA) (ThermoFisher) at 4°C and were subsequently stored at 4°C until imaged. The transgenic animals were imaged using a Zeiss LSM780 microscope.

### qRT-PCR analysis

Embryos were flash frozen on liquid nitrogen, approximately 150 embryos were used for each sample. RNA isolation was carried out following Invitrogen phenol-chloroform RNA extraction protocol. cDNA was then synthesized using iScript™ cDNA Synthesis Kit (BioRad). qRT-PCR reactions were prepared using SsoAdvanced Universal SYBR® Green Supermix (BioRad) and carried out on Real-Time PCR (CFX Opus 96) (BioRad).

Primers used:

18SF: CGG CTT AAT TTG ACT CAA CAC G

18SR: TTA GCA TGC CAG AGT CTC GTT C

Wnt4F: CGC CTA ACT ACT GCC ACA AA

Wnt4R: CCT CGC CCA CAA CAA AGA TA

SoxB1F: GTT GAC GGC TGA AGA GAA GG

SoxB1R: AGA ATT TGT CAA CCG CCA TC

LRRK2F: CCC ATA CCT CAC AGC TAC TTT AC

LRRK2R: CGA ATT CCG CCC TGT GTA TAA

### In situ hybridization

A 300bp fragment of the. NvLRRK2 coding region was cloned by PCR. The resulting PCR product was used to synthesize in situ probe by the addition of DIG-labeled UTP (Roche) plus the appropriate RNA Polymerase T7 or Sp6 (NEB). Probes were purified with RNA Clean Up Kit (Qiagen) and resuspended in 100uL of hybridization buffer.

*Nematostella* were collected at different time points after fertilization, fixed in 4%PFA overnight at 4C. Embryos or young larval animals were dehydrated in methanol series and stored at −20C. Before starting the *in-situ* protocol animals were rehydrated again through a methanol series back into phosphate buffered saline (PBS). In situs were carried out using published protocols [52].In summary, embryos were incubated in 1:1 PBST:Hyb for 30 minutes and pre-hybridized for 30 minutes. *NvLRRK2* probe was diluted into hybridization buffer and slides were allowed to hybridize overnight at 55°C. The following day the animals were washed 3 times in Wash Buffer (50% Formamide, 5x SSC and 0.1% Tween) for 30 minutes each, once in 1:1 Wash Buffer:PBST for 30 minutes and once in PBST at 55°C. Samples were rinsed in room temperature PBST 3 times for 5 minutes each before blocking buffer was added (2% goat serum, 2% BST in PBSTx) for 1 hour. Anti-DIG F_AB_ (Roche) was diluted 1:1000 in blocking buffer and slides were incubated for at least 1 hour. Samples were washed 3 times for 10 minutes each before addition of fresh AP Buffer for at least 10 minutes. Finally, samples were incubated in BM Purple (Roche) until colored reaction was observed. The reaction was stopped by several quick rinses in PBS and were fixed in 4% PFA for 10 minutes. Samples were mounted in 80% glycerol and images were taken with a Zeiss Discovery V8 microscope using Zen software.

### Statistical analysis

Statistical tests were performed using the Graphpad Prism 9 software. A D’Agostino and Pearson test was carried out to test for normality. Upon finding a non-normal distribution, a non-parametric Kruskal Wallis test was used to test for statistical significance between the medians of each measurement in the different experimental groups. A Dunn’s post-hoc multiple comparison test was used to compare the medians of the experimental groups against the control group

## Results

### *Nematostella vectenis* NvLRRK2 exhibits key residues conservation with human LRRK2

LRRK2 contains four protein-protein interaction domains, as well as domains conferring two distinct enzymatic activities. The kinase domain is a serine-threonine kinase capable of autophosphorylating residues within LRRK2, as well as heterologous range of substrates, most notably a subset of Rab GTPases [53–61].

The animalia display varying numbers of *LRRK* genes, ranging from 1 to 4 copies (figure 1A). In order to define the relationship between *Nematostella* nvLRRK2 and hsLRRK2, an evolutionary analysis into conservation of this gene was carried out. A phylogenetic tree using data from NCBI [44] and Clustal Omega [42, 43] was generated (figure 1B) to assess the relationship between representative LRRK genes across the animalia.

We then went on to compare the domain structure of *N. vectensis* and human LRRK2 to further determine the suitability of the *Nematostella* as a model for LRRK2-related PD (Figure 1C), using publicly available data sets from NCBI [44]. Consistent with conservation of LRRK proteins across evolution, these orthologues display a similar domain organisation, besides nvLRRK2 being a larger protein and lacking N-terminal armadillo repeats [62]. A Needleman-Wunsch algorithm was used to align the protein sequences to observe any biological similarities or differences [47]. This generated a score of 29% identity and 48% similarity across the open reading frame.

Parkinson’s Disease-causing mutations in hsLRRK2 are localized to the ROC, COR and kinase domains of the protein. As such, these domains in nvLRRK2 were compared to the human equivalent under the same global sequence alignment tool, the results of which are shown in Figure 1D. The scaffolding domains, Ankyrin and LRR, were also compared for a thorough comparison. The kinase domains were highly conserved, including the key kinase DYGI motif [63]. In contrast, the scaffolding and ROC/COR domains are less well conserved.

### Mutated residues are conserved in N. vectensis

To assess functional conservation across hsLRRK2 and nvLRRK2, residues that are mutated in human LRRK2 contributing to PD development were identified. These include R1441C/G, and N1437H (localized to the Roc domain), Y1699C (in the COR domain), G2019S and I2020T (in the kinase domain) [64–69]. Residues that are functionally significant and have been evolutionary conserved, are thought to have more of an impact in terms of disease status [70]. Figure 1E demonstrates that shows the most common PD associated LRRK2 mutation, G2019S, as well as the I2020T mutation residing in the same motif, is conserved in nvLRRK2.

Several of the amino acids surrounding these residues are also conserved, suggesting that these regions have not diverged throughout evolution despite selection pressures because of functional significance. The kinase domain appears to be the most well-conserved region of the protein, and the DYGI motif where these mutations occur are conserved across kinases, highlighting this as a critical region for kinase function [71]. In contrast, mutations in the ROC domain are less well-conserved, which mirrors the earlier analysis of a lower similarity score between nvLRRK2 and hsLRRK2 ROC and COR domains.

### Residues involved in post-translational modifications are not well conserved

To assess conservation of post-translational regulation between human and *Nematostella* LRRKs, residues previously reported to be involved in signal transduction were compared. Phosphorylation sites on LRRK2 are involved in regulating function and downstream signaling events [72]. Characterized phosphorylation sites in hsLRRK2 include S910, S935, S973, S955 located between the Ank and LRR domain, S1292 between LRR and Roc, and T2031, S2032 and T2035 in the kinase domain [72].

Unlike the mutated residues, the phosphorylated residues were not well conserved in both the LRR domain and the kinase domain. Figure 1F shows the sequence alignment demonstrating non-conserved residues as well as a conserved site. Only the S973 residue in the LRR domain and T2035 in the kinase domain were conserved in nvLRRK2, suggesting that the mechanisms involved in regulating signal transduction have diverged between these two proteins over evolution.

Scaffolding and structural domains in LRRK2 are less well conserved throughout species due to the lack of dependence for the functioning of the protein. Thus, is it unsurprising that these residues were not well conserved in nvLRRK2, and it suggests that post-translational modifications have perhaps drifted throughout evolution to suit higher or lower species depending on their functional requirements. This mirrors other studies into global phosphoproteomics that suggest phosphorylation events evolve individually between different species [73].

### NvLRRK2 kinase activity is necessary for faithful embryonic development

To investigate a role of the kinase activity of NvLrrk2 in regulating the development of *N.vectensis*, anemones were treated with structurally distinct validated inhibitors of LRRK2 kinase activity GNE-0877 and MLi-2. Freshly laid embryos were incubated in 5µM of respective inhibitor diluted in *Nematostella* water, while control embryos were grown in just water. The concentration of inhibitor to use was determined empirically by first testing concentrations in the range of 1-20µM, 5µM was chosen as for both inhibitors most of the embryos survived and showed a phenotype, while in higher concentrations a large percentage of the embryos died within a few days. Embryos incubated at 5µM developed through the first phases of development and reached the motile stage at relatively the same frequency as the control embryos. However, by 1 week post fertilization clear differences were observed, usually at this timepoint the embryos have developed tentacles and will commence feeding. Animals where the kinase activity of LRRK2 has been inhibited were overall smaller in length (Figure 2) and most showed no development of tentacles or stunted growth of tentacles (Figure 2) in comparison to controls that have much longer bodies and clear tentacles. Overall, the inhibitor treated animals display stunted inhibited growth from around 4 days post-fertilization onwards making it very difficult to determine if they developed normal organs like the pharynx and mesenteries.

**Figure 2.**
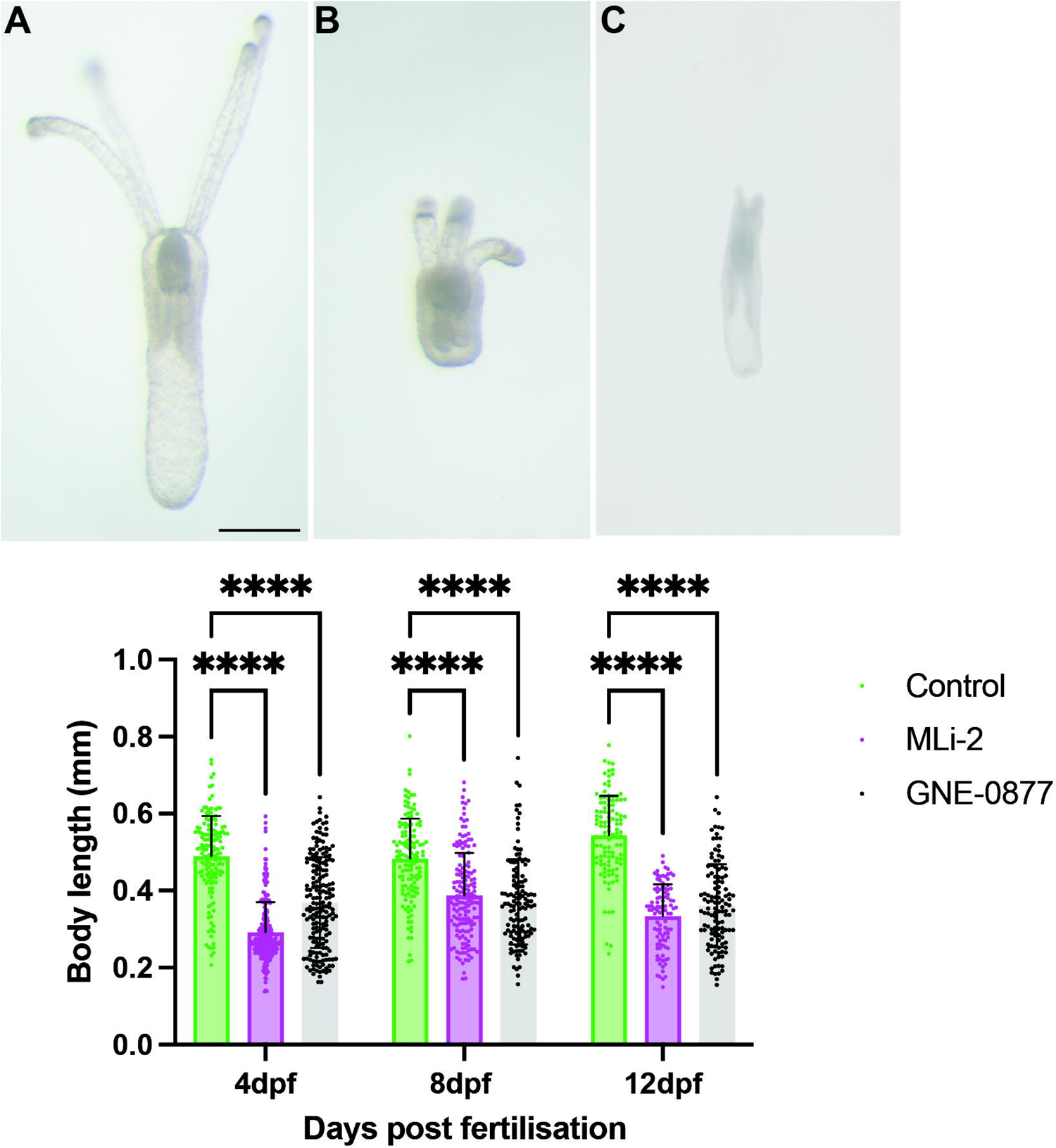
Pharmacologically inhibition of the NvLRRK2 kinase domain causes defects in embryonic development. (A) Control embryos 12 days post-fertilization, showing normal body length and development of internal organs like mesenteries and pharynx and external tentacles. (B-C) embryos grown in the presence of the Lrrk2 kinase inhibitor GNE-0877 or Mli-2 exhibit stunted body growth and failure to development proper tencles. Data shows mean +SD of body length in each groups; controls (12dpf n=120), GNE-0877 (12dpf n=121), and MLi-2 (12dpf n=130). Non-parametric post-hoc Dunn’s multiple comparison test following a significant Kruskal Wallis test.\**\*\*\*P≤0.0001, ***P≤ 0*.001, *\*\*P≤0.01, *P≤0.05,* ns=not significant. Scale bar = 1mm.

However, as all LRRK kinase inhibitor treated animals displayed stunted growth of tentacles or no tentacle growth we next examined expression of genes involved in specifying the oral region of the animal, *Wnt4*. *Wnt4* is expressed early in development from the planula stages and is important for induction of genes involved in specifying the oral region of the embryo [31, 74–77]. Here we find that at 48 hours post fertilization (hpf) in embryos exposed to the LRRK inhibitor that levels of *Wnt4* are significantly decreased in comparison to the control embryos, suggesting that the embryos are not inducing the genes necessary to specify head and to ultimately direct the embryos towards making tentacles (Figure 3A). As the tentacles are formed early in development and are highly innervated, we also examined induction and formation of neurons. *NvSoxB* is well-characterized to be expressed early in development in the cells that will form neurons and nematocysts [78, 79]. Quantitative analysis of *NvSoxB* levels in control embryos versus the LRRK2 kinase inhibitor treated animal discovered a significant lack of expression of *NvSoxB* in inhibitor treated animals (Figure 3B). We next examined the presence of development in the *Nematostella* embryos taking advantage of the NvLWamide-like::mCherry transgenic reporter line [18]. By one week post fertilization young animals have differentiated neurons with a complex network of axons in the body and tentacles as seen in Figure 3C. In comparison in animals exposed to the LRRK kinase inhibitor far fewer neurons are present, and they fail to extend axons and form networks (Figure 3D, E) suggesting that LRRK2 kinase activity is necessary for specification and proper differentiation of neurons during embryonic development. To determine if NvLRRK2 kinase activity is also necessary for maintenance of neurons in adult animals we placed 3 month and 6-month adult transgenic animals in LRRK2 kinase inhibitors and imaged their neurons after 4 days. We observed in all cases that the axons appeared to degenerate, and no axonal processes could be observed in the inhibitor treated animals versus control animals (Figure 2S), suggesting that the kinase activity of LRRK2 is necessary to maintain healthy connected axons

**Figure 3.**
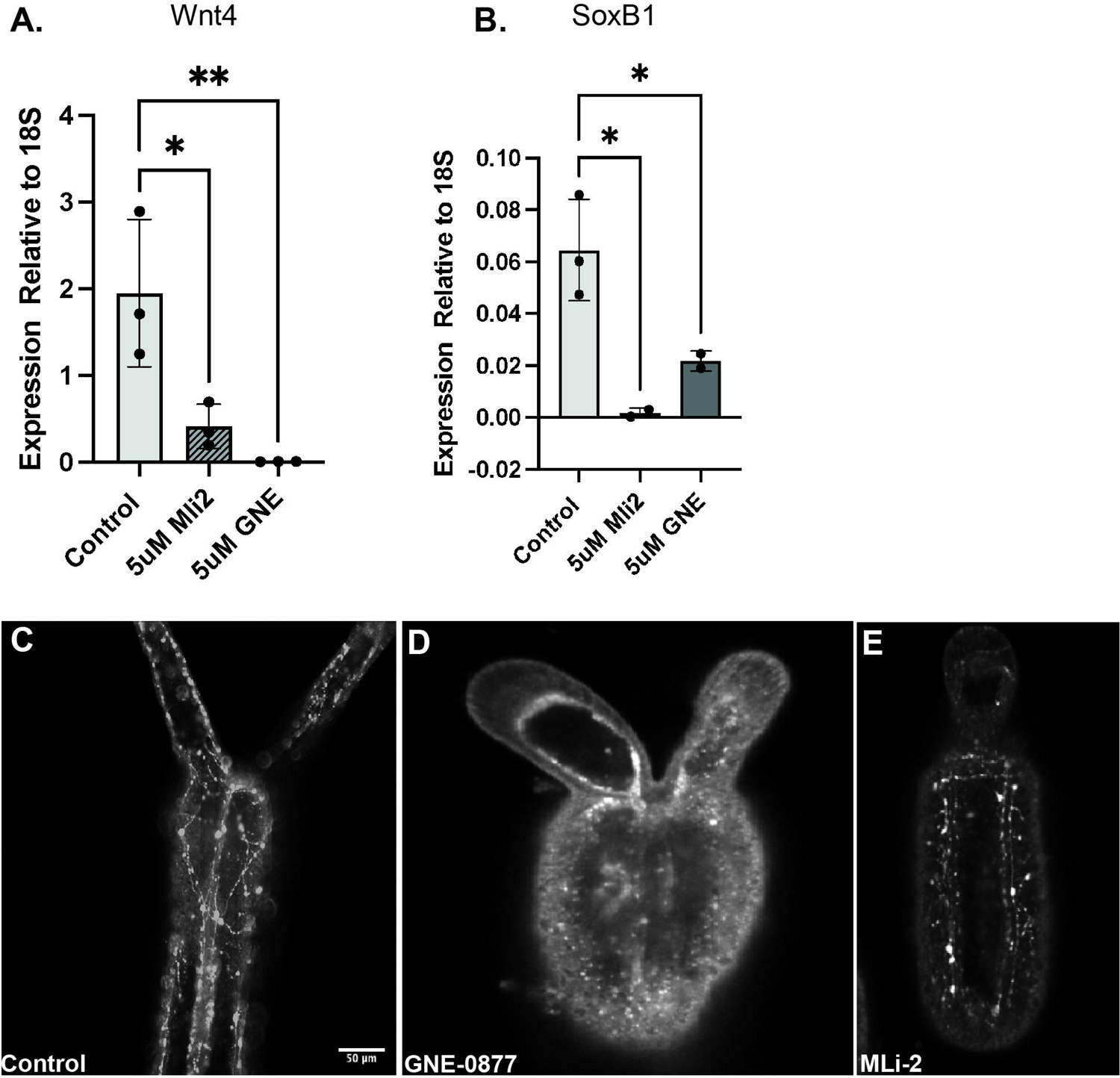
Inhibition of LRRK2 kinase activity leads to defects in neurons. (A) qRt-PCR of expression of the wnt4 gene that is necessary for induction of oral identity, levels are significantly decreased in inhibitor treated animals. (B) Quantification of the neuronal specification gene SoxB, kinase inhibition during embryonic development causes a decrease in SoxB expression levels. \**\*\*\*P≤0.0001, ***P≤ 0*.001, *\*\*P≤0.01*.(C-B) Confocal images of Nv-Lwamide-mcherry labelled neurons in developing embryos. By 7 days post fertilization a radial network of axons is visible in control embryos (C, n=150) while in Lrrk2 kinase treated embryos no network of axons is observed (D, E, n=160, n=140). Scale bar = 50µm.

### The LRRK2 kinase activity is necessary for regeneration in Nematostella

*Nematostella* are well-characterized to be capable of regeneration throughout life [28–30, 80–82]. The animal can be cut into multiple pieces and each fragment is capable of forming a whole new animal, the only exceptions to this is that a piece of foot alone or amputated tenacles are unable to facilitate full body regeneration. To assess whether LRRK2 has a role in regeneration in *N. vectensis*, the kinase activity of LRRK2 was inhibited after amputation below the pharynx. In adult animals’ pharynx and tentacle regeneration is complete within 7 days as seen here in control animals (Figure 4A). However, when exposed to the LRRK2 kinase inhibitor animals appeared to heal the wound but failed to regenerate significant tentacle, often both the pharynx and tentacles were missing. (Figure 4). Additionally, there was a difference in body length for those treated with the kinase inhibitor, suggesting that like in the homeostatic conditions lack of a functional LRRK2 kinase leads to degeneration of the axons and overall shrinking of the body axis (Figure 2S). Taken together these data suggest that nvLRRK2 plays an important role both in the development, homeostasis, and regeneration of neurons.

**Figure 4.**
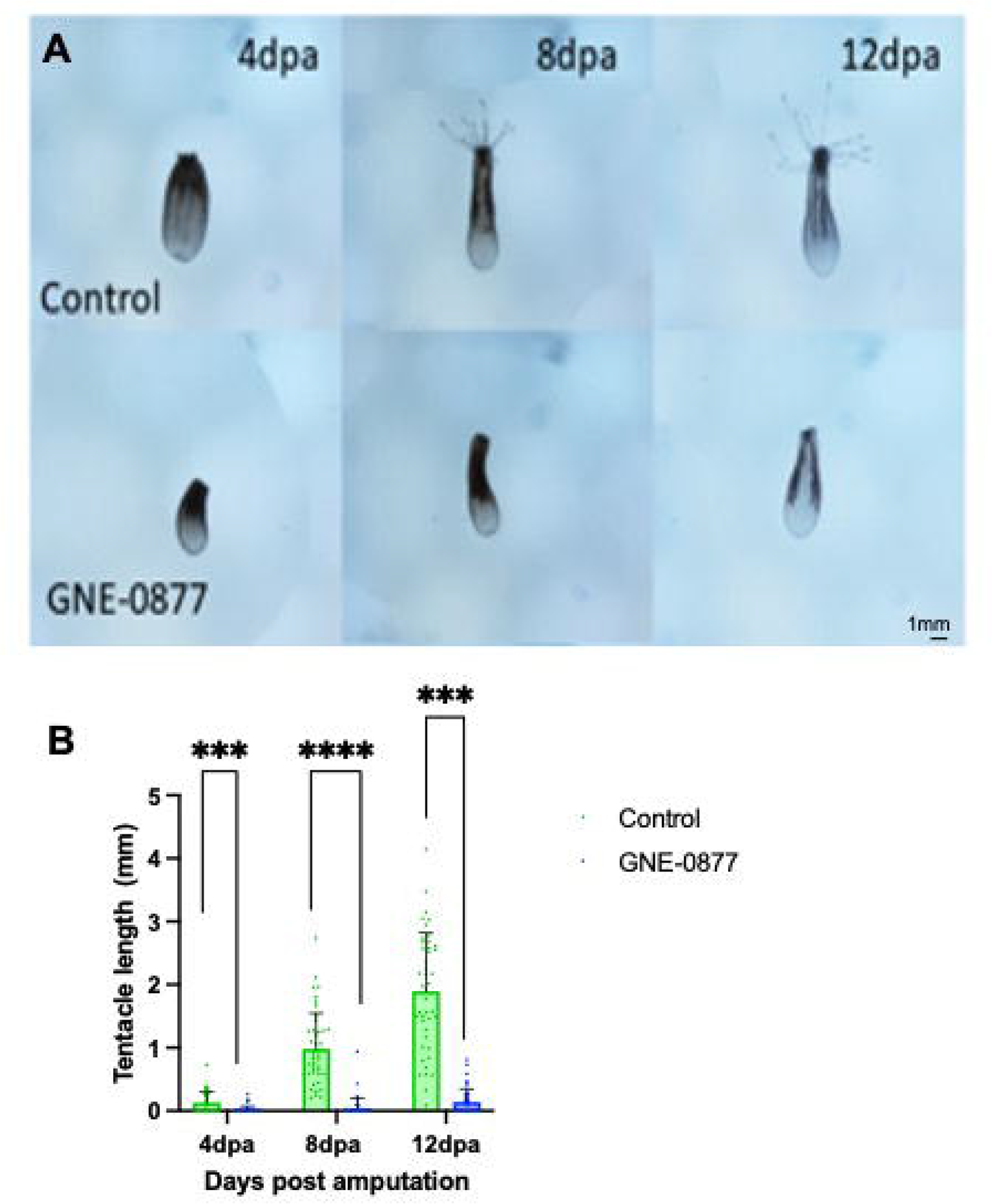
Pharmacological LRRK2 kinase inhibition impacts regeneration of *N. vec*tensis tentacles. (A) Control animals regenerate their oral portions including tentacles withing 12 days in adult animals, in comparison animals exposed to the Lrrk2 kinase inhibitor GNE-0877 fail to regenerate tentacles. (B) Data shows mean + SD of tentacle length in each of the experimental groups; controls (4dpa n=44, 8dpa n=45, 12dpa n=44), GNE-0877 (4dpa n=45, 8dpa n=44, 12dpa n=44). Non-parametric Dunn’s multiple comparison test following a significant Kruskal Wallis. ns = not significant, *\*\*\*\*P≤0.0001, ***P≤0.001, **P≤0.01 *P≤0.05*.

## Discussion

In this study, we have introduced a novel organism for studying LRRK biology, and demonstrated that its LRRK2 orthologue, nvLRRK2, has great similarity to hsLRRK2. It has previously been noted that *Nematostella* are more similar to vertebrates on a genomic scale, than more well-used organisms such as the *Drosophila* are [75]. This study expands on this and demonstrates its great potential for providing information regarding the function of LRRK2 in a disease context.

Here we have taken advantage of the availability of large number of *Nematostella* embryos and of commercially available validated LRRK2 kinase inhibitors to determine the function of nvLRRK2. We have uncovered a key role for the kinase activity of this gene in promoting normal embryonic growth, in specification of the oral region of the animal and in neuronal differentiation. Additionally, we have demonstrated that the kinase activity of LRRK2 is necessary to maintain a functional radial network of nerves in the *Nematostella* and also to promote head regeneration where a complex set of nerves must be regenerated. This study indicates conservation of a gene which has been primarily studied in vertebrates and illustrates its high conservation in invertebrates. In previous studies, LRRK2 has been identified as making an important contribution to regeneration in a large-scale transcriptional profiling approach during planaria regeneration, where the authors suggest it may play a role in activated of the neoblast populations that are essential for planaria regeneration [40]. It will be important to determine in the future the identity of the effector pathway used by nvLRRK2 to regulate regeneration and development, providing mechanistic insight into this process. In mammals it is well-established that a subset of Rab proteins – including Rab10 - are LRRK2 substrates [83], undergoing phosphorylation by LRRK2 [84]. We were able to confirm that *Nematostella* has a Rab10 ortholog, but in the absence of specific tool antibodies to investigate this were unable to test for direct phosphorylation by nvLRRK2. Likewise, the conservation of key residues that are mutated in human disease (G2019S and I2020T) with defined biochemical consequences provides an opportunity to assess the implications of gain of LRRK2 function on regeneration and development in the Nematostella system.

Our results open up a new avenue of investigation into the physiological role and importance of *LRRK* genes and provides a novel model platform to test the consequences of modulating LRRK2 function – with the potential to provide insights relevant to human disease.

## List of Abbreviations

Dpf: days post fertilization

Dpa: days post amputation

LRRK: leucine rich repeat kinase

NvLRRK2: *Nematostella vectensis* leucine rich repeat kinase 2

PD: Parkinsons disease

PBS: Phosphate buffered saline

PFA: paraformaldehyde

## Acknowledgements

G.H. was supported by a travel grant from the Company of Biologists. The collaboration between P.A.L. and K.E. is supported by Royal Society International Exchange Grant IES\R3\203078. KE is supported by a grant from NICHD R01 HD092451, start-up funds from the MBL and funding from the Owens Family Foundation. P.A.L. is a Royal Society Industry Fellow (fellowship IF\R2\222002). This research was funded in whole or in part by Aligning Science Across Parkinson’s (grant number ASAP 000478) through the Michael J. Fox Foundation for Parkinson’s Research (MJFF). For the purpose of open access, the author has applied a CC BY public copyright licence to all Author Accepted Manuscripts arising from this submission.

---

**Figure S1.** (A-C) *In sit*u hybridization of *NvLrrk2* during Nematostella embryonic development. (D) Quantification real time PCR analysis of Nematostella *NvLrrk2* gene during the first week of development in Nematostella embryos.

**Figure S2.** LRRK2 kinase inhibition results in neuronal degeneration in adult Nematostella. NvLWamide-mCherry neurons imaged in control animals, nerve networks are clearly visible (A), after 4 days exposure to kinase inhibitors GNE-0887 (B) or Mli-2 (C) very few axons are observed in adult Nematostella, axons appear to be degenerating. (control N=60, GNE-0887 n=52, Mli-2 N=45). Scale bar = 75µm.

## References

1. Layden, M.J., F. Rentzsch, and E. Rottinger, The rise of the starlet sea anemone Nematostella vectensis as a model system to investigate development and regeneration. Wiley Interdiscip Rev Dev Biol, 2016. 5(4): p. 408–28.

2. Miller, D.J., E.E. Ball, and U. Technau, Cnidarians and ancestral genetic complexity in the animal kingdom. Trends Genet, 2005. 21(10): p. 536–9.

3. Warner, J.F., et al., NvERTx: a gene expression database to compare embryogenesis and regeneration in the sea anemone Nematostella vectensis. Development, 2018. 145(10).

4. Sebe-Pedros, A., et al., Cnidarian Cell Type Diversity and Regulation Revealed by Whole-Organism Single-Cell RNA-Seq. Cell, 2018. 173(6): p. 1520–1534.e20.

5. Ikmi, A., et al., TALEN and CRISPR/Cas9-mediated genome editing in the early-branching metazoan Nematostella vectensis. Nat Commun, 2014. 5: p. 5486.

6. Desvignes, T., P. Pontarotti, and J. Bobe, Nme gene family evolutionary history reveals pre-metazoan origins and high conservation between humans and the sea anemone, Nematostella vectensis. PLoS One, 2010. 5(11): p. e15506.

7. Putnam, N.H., et al., Sea anemone genome reveals ancestral eumetazoan gene repertoire and genomic organization. Science, 2007. 317(5834): p. 86–94.

8. Steger, J., et al., Single-cell transcriptomics identifies conserved regulators of neuroglandular lineages. Cell Rep, 2022. 40(12): p. 111370.

9. Zimmermann, B., et al., Sea anemone genomes reveal ancestral metazoan chromosomal macrosynteny. bioRxiv, 2020.

10. Marin, I., Ancient origin of the Parkinson disease gene LRRK2. J Mol Evol, 2008. 67(1): p. 41–50.

11. Rentzsch, F., et al., FGF signalling controls formation of the apical sensory organ in the cnidarian Nematostella vectensis. Development, 2008. 135(10): p. 1761–9.

12. Amiel, A.R., et al., A bipolar role of the transcription factor ERG for cnidarian germ layer formation and apical domain patterning. Dev Biol, 2017. 430(2): p. 346–361.

13. Technau, U., Gastrulation and germ layer formation in the sea anemone Nematostella vectensis and other cnidarians. Mech Dev, 2020. 163: p. 103628.

14. Salinas-Saavedra, M., A.Q. Rock, and M.Q. Martindale, Germ layer-specific regulation of cell polarity and adhesion gives insight into the evolution of mesoderm. Elife, 2018. 7.

15. Nakanishi, N. and M.Q. Martindale, CRISPR knockouts reveal an endogenous role for ancient neuropeptides in regulating developmental timing in a sea anemone. Elife, 2018. 7.

16. Nakanishi, N., et al., Nervous systems of the sea anemone Nematostella vectensis are generated by ectoderm and endoderm and shaped by distinct mechanisms. Development, 2012. 139(2): p. 347–57.

17. Havrilak, J.A., et al., Characterization of the dynamics and variability of neuronal subtype responses during growth, degrowth, and regeneration of Nematostella vectensis. BMC Biol, 2021. 19(1): p. 104.

18. Havrilak, J.A., et al., Characterization of NvLWamide-like neurons reveals stereotypy in Nematostella nerve net development. Dev Biol, 2017. 431(2): p. 336–346.

19. Bylund, M., et al., Vertebrate neurogenesis is counteracted by Sox1-3 activity. Nat Neurosci, 2003. 6(11): p. 1162–8.

20. Graham, V., et al., SOX2 functions to maintain neural progenitor identity. Neuron, 2003. 39(5): p. 749–65.

21. Sandberg, M., M. Kallstrom, and J. Muhr, Sox21 promotes the progression of vertebrate neurogenesis. Nat Neurosci, 2005. 8(8): p. 995–1001.

22. Galliot, B., et al., Origins of neurogenesis, a cnidarian view. Dev Biol, 2009. 332(1): p. 2–24.

23. Watanabe, H., et al., Sequential actions of beta-catenin and Bmp pattern the oral nerve net in Nematostella vectensis. Nat Commun, 2014. 5: p. 5536.

24. Sebe-Pedros, A., et al., Cnidarian Cell Type Diversity and Regulation Revealed by Whole-Organism Single-Cell RNA-Seq. Cell, 2018. 173(6): p. 1520–1534 e20.

25. Jegla, T., et al., Expanded functional diversity of shaker K(+) channels in cnidarians is driven by gene expansion. PLoS One, 2012. 7(12): p. e51366.

26. Babonis, L.S. and M.Q. Martindale, Old cell, new trick? Cnidocytes as a model for the evolution of novelty. Integr Comp Biol, 2014. 54(4): p. 714–22.

27. DuBuc, T.Q., N. Traylor-Knowles, and M.Q. Martindale, Initiating a regenerative response; cellular and molecular features of wound healing in the cnidarian Nematostella vectensis. BMC Biol, 2014. 12: p. 24.

28. Amiel, A.R., et al., Characterization of Morphological and Cellular Events Underlying Oral Regeneration in the Sea Anemone, Nematostella vectensis. Int J Mol Sci, 2015. 16(12): p. 28449–71.

29. Amiel, A.R., et al., [The sea anemone Nematostella vectensis, an emerging model for biomedical research: Mechano-sensitivity, extreme regeneration and longevity]. Med Sci (Paris), 2021. 37(2): p. 167–177.

30. Amiel, A.R. and E. Röttinger, Experimental Tools to Study Regeneration in the Sea Anemone Nematostella vectensis. Methods Mol Biol, 2021. 2219: p. 69–80.

31. Schaffer, A.A., et al., A transcriptional time-course analysis of oral vs. aboral whole-body regeneration in the Sea anemone Nematostella vectensis. BMC Genomics, 2016. 17(1): p. 718.

32. Zhu, C., S. Herbst, and P.A. Lewis, Leucine-rich repeat kinase 2 at a glance. J Cell Sci, 2023. 136(17).

33. Schlitter, A.M., et al., The LRRK2 gene in Parkinson’s disease: mutation screening in patients from Germany. J Neurol Neurosurg Psychiatry, 2006. 77(7): p. 891–2.

34. Berwick, D.C. and K. Harvey, LRRK2: an eminence grise of Wnt-mediated neurogenesis? Front Cell Neurosci, 2013. 7: p. 82.

35. Myasnikov, A., et al., Structural analysis of the full-length human LRRK2. Cell, 2021. 184(13): p. 3519–3527 e10.

36. van Egmond, W.N. and P.J. van Haastert, Characterization of the Roco protein family in Dictyostelium discoideum. Eukaryot Cell, 2010. 9(5): p. 751–61.

37. Wauters, L., W. Versees, and A. Kortholt, Roco Proteins: GTPases with a Baroque Structure and Mechanism. Int J Mol Sci, 2019. 20(1).

38. Alessi, D.R. and E. Sammler, LRRK2 kinase in Parkinson’s disease. Science, 2018. 360(6384): p. 36–37.

39. Lewis, P.A., A step forward for LRRK2 inhibitors in Parkinson’s disease. Sci Transl Med, 2022. 14(648): p. eabq7374.

40. Benham-Pyle, B.W., et al., Identification of rare, transient post-mitotic cell states that are induced by injury and required for whole-body regeneration in Schmidtea mediterranea. Nat Cell Biol, 2021. 23(9): p. 939–952.

41. Klein, S., et al., Common Environmental Pollutants Negatively Affect Development and Regeneration in the Sea Anemone Nematostella vectensis Holobiont. Frontiers in Ecological Evolution 2021. 9.

42. Goujon, M., et al., A new bioinformatics analysis tools framework at EMBL-EBI. Nucleic Acids Res, 2010. 38(Web Server issue): p. W695-9.

43. Sievers, F., et al., Fast, scalable generation of high-quality protein multiple sequence alignments using Clustal Omega. Mol Syst Biol, 2011. 7: p. 539.

44. Sayers, E.W., et al., Database resources of the national center for biotechnology information. Nucleic Acids Res, 2022. 50(D1): p. D20–D26.

45. Altschul, S.F., et al., Basic local alignment search tool. J Mol Biol, 1990. 215(3): p. 403–10.

46. Altschul, S.F., et al., Gapped BLAST and PSI-BLAST: a new generation of protein database search programs. Nucleic Acids Res, 1997. 25(17): p. 3389–402.

47. Needleman, S.B. and C.D. Wunsch, A general method applicable to the search for similarities in the amino acid sequence of two proteins. J Mol Biol, 1970. 48(3): p. 443–53.

48. Jararweh, Y., et al., Improving the performance of the needleman-wunsch algorithm using parallelization and vectorization techniques. Multimedia Tools and Applications 2017. 78: p. 3961–3977.

49. Cookson, M.R., The role of leucine-rich repeat kinase 2 (LRRK2) in Parkinson’s disease. Nat Rev Neurosci, 2010. 11(12): p. 791–7.

50. Mills, R.D., et al., Analysis of LRRK2 accessory repeat domains: prediction of repeat length, number and sites of Parkinson’s disease mutations. Biochem Soc Trans, 2012. 40(5): p. 1086–9.

51. Rui, Q., et al., The Role of LRRK2 in Neurodegeneration of Parkinson Disease. Curr Neuropharmacol, 2018. 16(9): p. 1348–1357.

52. Wolenski, F.S., et al., Characterizing the spatiotemporal expression of RNAs and proteins in the starlet sea anemone, Nematostella vectensis. Nat Protoc, 2013. 8(5): p. 900–15.

53. Baptista, M.A.S., et al., LRRK2 inhibitors induce reversible changes in nonhuman primate lungs without measurable pulmonary deficits. Sci Transl Med, 2020. 12(540).

54. Dodson, M.W., et al., Roles of the Drosophila LRRK2 homolog in Rab7-dependent lysosomal positioning. Hum Mol Genet, 2012. 21(6): p. 1350–63.

55. Eguchi, T., et al., LRRK2 and its substrate Rab GTPases are sequentially targeted onto stressed lysosomes and maintain their homeostasis. Proc Natl Acad Sci U S A, 2018. 115(39): p. E9115–E9124.

56. Fdez, E., et al., Pathogenic LRRK2 regulates centrosome cohesion via Rab10/RILPL1-mediated CDK5RAP2 displacement. iScience, 2022. 25(6): p. 104476.

57. Huber, L.A., et al., Rab8, a small GTPase involved in vesicular traffic between the TGN and the basolateral plasma membrane. J Cell Biol, 1993. 123(1): p. 35–45.

58. Lara Ordonez, A.J., et al., RAB8, RAB10 and RILPL1 contribute to both LRRK2 kinase-mediated centrosomal cohesion and ciliogenesis deficits. Hum Mol Genet, 2019. 28(21): p. 3552–3568.

59. Lis, P., et al., Development of phospho-specific Rab protein antibodies to monitor in vivo activity of the LRRK2 Parkinson’s disease kinase. Biochem J, 2018. 475(1): p. 1–22.

60. MacLeod, D.A., et al., RAB7L1 interacts with LRRK2 to modify intraneuronal protein sorting and Parkinson’s disease risk. Neuron, 2013. 77(3): p. 425–39.

61. Malik, A.U., et al., Deciphering the LRRK code: LRRK1 and LRRK2 phosphorylate distinct Rab proteins and are regulated by diverse mechanisms. Biochem J, 2021. 478(3): p. 553–578.

62. Marin, I., W.N. van Egmond, and P.J. van Haastert, The Roco protein family: a functional perspective. FASEB J, 2008. 22(9): p. 3103–10.

63. Schmidt, S.H., et al., Conformation and dynamics of the kinase domain drive subcellular location and activation of LRRK2. Proc Natl Acad Sci U S A, 2021. 118(23).

64. Funayama, M., et al., A new locus for Parkinson’s disease (PARK8) maps to chromosome 12p11.2-q13.1. Ann Neurol, 2002. 51(3): p. 296–301.

65. Goldwurm, S., et al., The G6055A (G2019S) mutation in LRRK2 is frequent in both early and late onset Parkinson’s disease and originates from a common ancestor. J Med Genet, 2005. 42(11): p. e65.

66. Greene, I.D., et al., Evidence that the LRRK2 ROC domain Parkinson’s disease-associated mutants A1442P and R1441C exhibit increased intracellular degradation. J Neurosci Res, 2014. 92(4): p. 506–16.

67. Kalia, L.V., et al., Clinical correlations with Lewy body pathology in LRRK2-related Parkinson disease. JAMA Neurol, 2015. 72(1): p. 100–5.

68. Puschmann, A., et al., First neuropathological description of a patient with Parkinson’s disease and LRRK2 p.N1437H mutation. Parkinsonism Relat Disord, 2012. 18(4): p. 332–8.

69. Ruiz-Martinez, J., et al., Penetrance in Parkinson’s disease related to the LRRK2 R1441G mutation in the Basque country (Spain). Mov Disord, 2010. 25(14): p. 2340–5.

70. Kim, D., et al., Evolutionary coupling analysis identifies the impact of disease-associated variants at less-conserved sites. Nucleic Acids Res, 2019. 47(16): p. e94.

71. DiBona, D.R., Functional analysis of tight junction organization. Pflugers Arch, 1985. 405 **Suppl 1**: p. S59–66.

72. Price, A., et al., The LRRK2 signalling system. Cell Tissue Res, 2018. 373(1): p. 39–50.

73. Pasquier, C. and A. Robichon, Evolutionary Divergence of Phosphorylation to Regulate Interactive Protein Networks in Lower and Higher Species. Int J Mol Sci, 2022. 23(22).

74. Wijesena, N., D.K. Simmons, and M.Q. Martindale, Antagonistic BMP-cWNT signaling in the cnidarian Nematostella vectensis reveals insight into the evolution of mesoderm. Proc Natl Acad Sci U S A, 2017. 114(28): p. E5608–e5615.

75. Sullivan, J.C., et al., Conserved and novel Wnt clusters in the basal eumetazoan Nematostella vectensis. Development Genes and Evolution, 2007. 217(3): p. 235–239.

76. Lee, P.N., et al., A WNT of things to come: evolution of Wnt signaling and polarity in cnidarians. Semin Cell Dev Biol, 2006. 17(2): p. 157–67.

77. DuBuc, T.Q., et al., Hox and Wnt pattern the primary body axis of an anthozoan cnidarian before gastrulation. Nat Commun, 2018. 9(1): p. 2007.

78. Magie, C.R., K. Pang, and M.Q. Martindale, Genomic inventory and expression of Sox and Fox genes in the cnidarian Nematostella vectensis. Dev Genes Evol, 2005. 215(12): p. 618–30.

79. Richards, G.S. and F. Rentzsch, Transgenic analysis of a SoxB gene reveals neural progenitor cells in the cnidarian Nematostella vectensis. Development, 2014. 141(24): p. 4681–9.

80. Bossert, P. and G.H. Thomsen, Inducing Complete Polyp Regeneration from the Aboral Physa of the Starlet Sea Anemone Nematostella vectensis. J Vis Exp, 2017(119).

81. Bossert, P.E., M.P. Dunn, and G.H. Thomsen, A staging system for the regeneration of a polyp from the aboral physa of the anthozoan Cnidarian Nematostella vectensis. Dev Dyn, 2013. 242(11): p. 1320–31.

82. Schaffer, A.A., et al., A transcriptional time-course analysis of oral vs. aboral whole-body regeneration in the Sea anemone Nematostella vectensis. BMC Genomics, 2016. 17: p. 718.

83. Steger, M., et al., Phosphoproteomics reveals that Parkinson’s disease kinase LRRK2 regulates a subset of Rab GTPases. Elife, 2016. 5.

84. Ordóñez, A., et al., LRRK2 causes centrosomal deficits via phosphorylated Rab10 and RILPL1 at centriolar subdistal appendages. BioRxiv, 2021.

