## Supplementary figures and images for "LRRK2 kinase activity is necessary for development and regeneration in *Nematostella vectensis*"

### Supplementary Figure 1

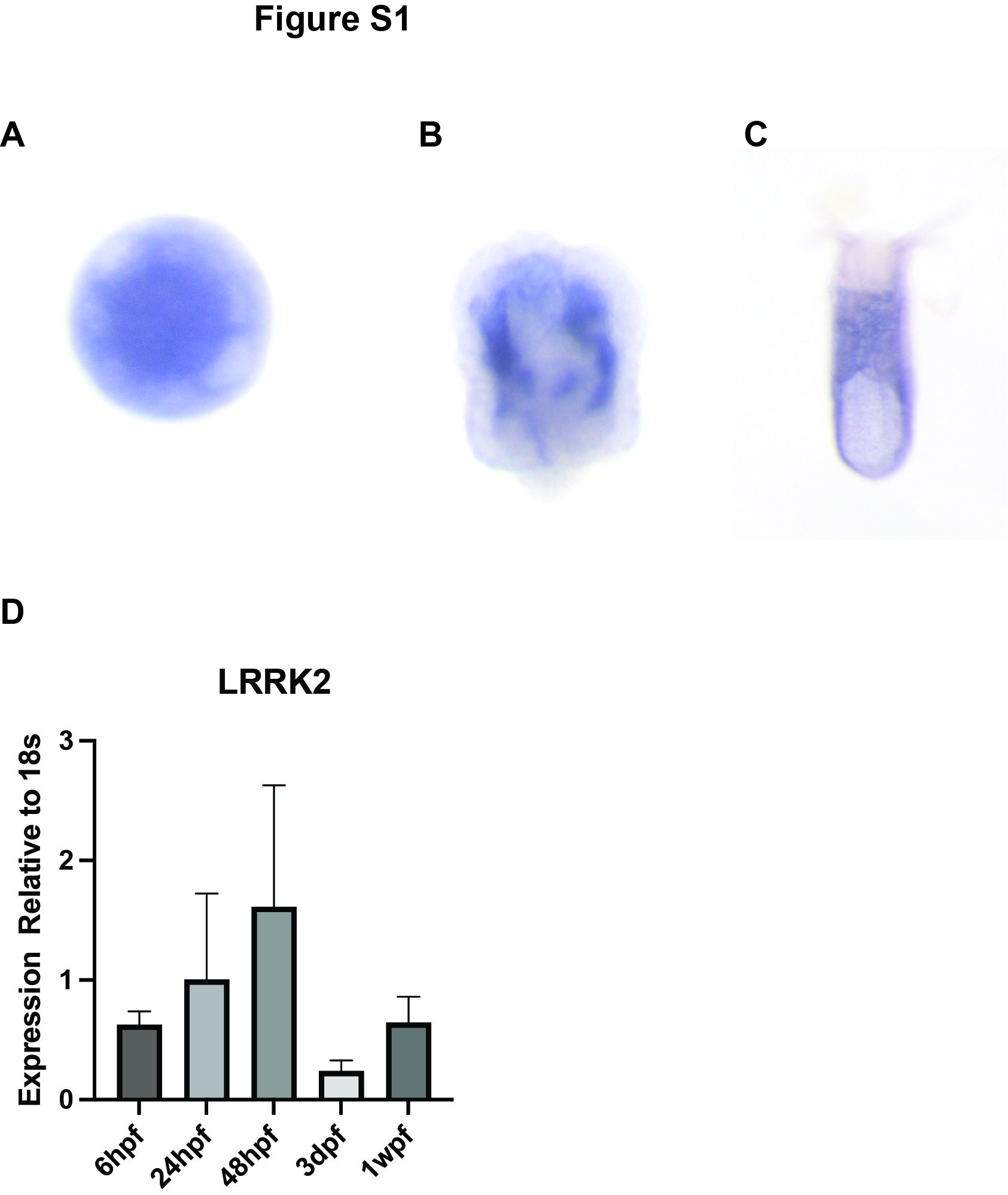

### Supplementary Figure 2

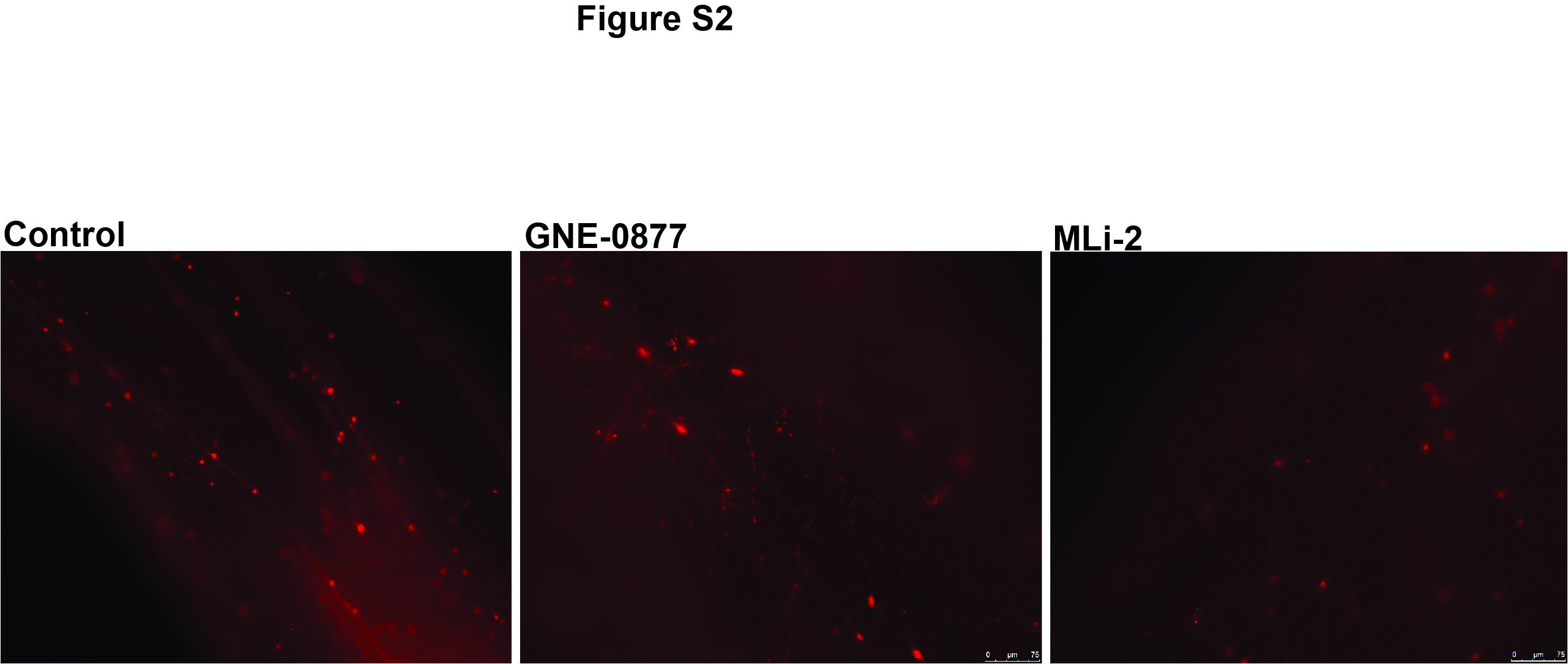
